# The relationship between the spatial scaling of biodiversity and ecosystem stability

**DOI:** 10.1101/162511

**Authors:** Robin Delsol, Michel Loreau, Bart Haegeman

**Affiliations:** Centre for Biodiversity Theory and Modelling, Theoretical and Experimental Ecology Station, CNRS and Paul Sabatier University, Moulis, France

## Abstract

Ecosystem stability and its link with biodiversity have been mainly studied at the local scale. Here we present a simple theoretical model to address the joint dependence of diversity and stability on spatial scale, from local to continental. The notion of stability we use is based on the temporal variability of an ecosystem-level property, such as primary productivity. In this way, our model integrates the well-known species-area relationship (SAR) with a recent proposal to quantify the spatial scaling of stability, called the invariability-area relationship (IAR). To explore the possible links between the two relationships, we contrast two assumptions about the spatial decay of correlations. In case species differences determine spatial decorrelation, the IAR is a duplicate of the SAR; in case decorrelation is directly determined by spatial separation, the IAR is unrelated to the SAR. We apply the two model variants to explore the effects of species loss and habitat destruction on stability, and find a rich variety of multi-scale spatial dependencies. Our work emphasizes the importance of studying diversity and stability across spatial scales, and provides a point of reference for mechanistic models and data analyses.

## Introduction

Decades of ecological research have explored how features of ecosystems affect their stability (May, 1973; Pimm, 1984; McCann, 2000; Ives & Carpenter, 2007). Because of the worldwide loss of biodiversity, most studies on ecosystem stability have focused on the role played by biodiversity, resulting in an extensive debate on the diversity-stability relationship. Although this debate is not yet settled, there is now strong empirical evidence and theoretical support that biodiversity tends to increase the stability of ecosystem processes (Lehman & Tilman, 2000; Tilman *et al.*, 2006; Jiang & Pu, 2009; Campbell *et al.*, 2011; Loreau & de Mazancourt, 2013; Gross *et al.*, 2014).

To date, the majority of studies addressing the effects of biodiversity on ecosystem stability have dealt with small spatial scales, such as microcosms (Petchey *et al.*, 2002; Steiner *et al.*, 2005) and grassland experiments (Bai *et al.*, 2004; Tilman *et al.*, 2006). Yet sustaining ecosystem structure, functioning and services requires a broader understanding of stability across a wide range of scales. Therefore there is a current need to better understand how biodiversity regulates ecosystem stability at larger spatial scales that are more relevant to ecosystem management (Peterson *et al.*, 1998; Chalcraft, 2013; Isbell *et al.*, 2017).

The spatial scaling of biodiversity is one of the most studied ecological patterns (Rosenzweig, 1995). In particular, the species-area relationship (SAR) describes how species richness *S* changes with area *A*. When increasing the observation area, additional species can be observed, so that the SAR is increasing. Typically, for a limited range of intermediate spatial scales, empirical SARs are well approximated by a power-law function, *S* = *cA*^*z*^ where *c* and *z* are empirical constants. When very small and very large scales are also included, SARs often exhibit three distinct phases on a log-log plot: concave at local scales, approximately linear at regional scales, and convex at continental scales (Rosenzweig, 1995; Hubbell, 2001; Storch *et al.*, 2012; see also Fig. 1). Various simple models have been proposed to explain this triphasic shape (Chave *et al.*, 2002; Allen & White, 2003; Rosindell & Cornell, 2007; Harte *et al.*, 2009).

**Figure 1.**
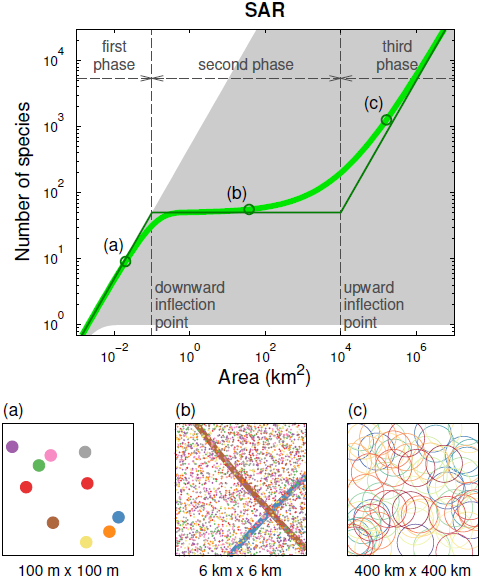
Our model predicts a triphasic species-area relationship (SAR). A log-log plot of number of species *S*(*A*) against observation area *A* has three distinct phases. Top panel: exact solution of model (thick green line) and piecewise linear approximation (thin green line). The shaded region indicates the set of possible SARs for a fixed configuration of individuals. For points on the upper boundary, each individual belongs to a different species (see also Storch 2016); for points on the lower boundary, all individuals belong to the same species. Bottom panel (a): In the first phase almost all individuals (represented by dots) in the observation area belong to different species (represented by colors). Bottom panel (b): In the second phase the observed species have many individuals in the observation area. Some species range boundaries (represented by lines) are visible in the observation area. For clarity, only 20 percent of the species are shown. Bottom panel (c): In the third phase, the species ranges of the observed species (represented by circles) are for the most part included in the observed area. For clarity, only 10 percent of the species are shown. Parameter values: *λ*_*I*_ = 10, *λ*_*S*_ = 0.005, *Q* = 10^4^.

In contrast, the spatial scaling of ecosystem stability has hardly been studied. One problem is that ecological stability is a multi-faceted concept, for which numerous measures have been proposed (Pimm, 1984; Grimm & Wissel, 1997). For many of them it is unclear how they can be scaled up to larger spatial scales. An exception are stability measures based on temporal variability (Wang & Loreau, 2014). Indeed, the temporal variability of an ecosystem property such as total biomass or productivity can be readily quantified for areas of different size. Using invariability as a measure of stability, Wang *et al.* (2017) proposed the invariability-area relationship (IAR) to describe the spatial scaling of ecosystem stability. They showed that, similarly to SARs, empirical IARs have a triphasic shape on a log-log plot, suggesting a connection between SARs and IARs.

Variability-based stability, which is commonly used in empirical studies (Jiang & Pu, 2009; Donohue *et al.*, 2016), is strongly determined by temporal correlation or synchrony among species or across space (Wang & Loreau, 2014). For example, consider the total invariability of two spatially separated parts of an ecosystem. If the temporal fluctuations of the parts are perfectly correlated, the total invariability is equal to (an appropriate average of) the invariability of the parts. The more decorrelated or asynchronous the fluctuations, the larger the difference between the total invariability and the invariability of the parts, that is, the larger the stability gain. As a consequence, the IAR is increasing, and its rate of increase, i.e., its slope, at a particular spatial scale is governed by the correlation between parts of the ecosystem at that scale.

Hence, the spatial scaling of invariability can be understood in terms of temporal correlation at different spatial scales. In this paper we consider two very different assumptions about these correlations. First, we assume that species differences are the main source of spatial asynchrony. Because different parts of the ecosystem are populated by different species, this leads to decorrelation between the parts, and hence increased total invariability. This assumption is related to statistical explanations of the local diversity-stability relationship, such as the portfolio effect and the insurance hypothesis (May, 1974; Tilman *et al.*, 1998; Yachi & Loreau, 1999; Thibaut & Connolly, 2013). Second, we assume that spatial distance is at the origin of asynchrony. This can occur when the correlations of the environmental disturbances determine the correlations of the ecosystem fluctuations directly, that is, without being mediated by species differences. This mechanism is related to the Moran effect in population ecology (Hudson & Cattadori, 1999; Liebhold *et al.*, 2004). Clearly, under the second assumption we expect a weaker relationship between diversity and stability than under the first one.

We start by introducing a minimal model that incorporates the SAR and the IAR, and use it to clarify their relationship. The model predicts triphasic curves for both the SAR and the IAR, in qualitative agreement with empirical data. Then, we implement the two correlation assumptions. Under the first assumption, which we call decorrelation by species turnover (DST), the IAR essentially coincides with the SAR. Under the second assumption, called decorrelation by distance (DD), the IAR is generally unrelated to the SAR. Nevertheless, there is a range of parameter values for which the IAR-DD closely resembles the IAR-DST. Next, we subject the two model variants to different scenarios of species loss and habitat destruction, and show that the response of stability across spatial scales differs markedly between the two correlation assumptions. We conclude by discussing the implications of these findings, and argue that our simple model could serve as a framework for more mechanistic models.

## Methods

### Modelling approach

We construct a minimal model that simultaneously predicts the SAR and the IAR. To predict the SAR, we specify the geographical ranges of the species. Indeed, if species ranges are given, one can readily determine how many species are present in any specific area. While mechanistic models determine these ranges based on more detailed ecological variables, such as habitat preferences, dispersal properties and interaction strengths (e.g., Rybicki & Hanski 2013; Matias *et al.* 2014), here we consider species ranges as a model input. To predict the IAR, we specify how the entities making up the ecosystem fluctuate through time. As detailed below, the intensities and spatial correlations of these fluctuations suffice to determine invariability in any specific area. Again, mechanistic models could explain these fluctuation characteristics in terms of ecological processes, such as species interactions, dispersal and responses to environmental disturbances. In our model, however, the statistical properties of these fluctuations are an assumption, not a prediction.

While this model setup is rather general, for ease of presentation we discuss it in the context of a more specific system. We propose to look at the species diversity of plants and the variability of their productivity. The productivity of individual plants varies from year to year, and these individual-level fluctuations add up to generate variability of the primary productivity at the ecosystem level. The latter, together with the spatial distribution of individuals and species, determine the invariability-area relationship. We now describe the model. Variables and parameters are summarized in Table 1. Mathematical details are presented in the appendices.

**Table 1.** List of model variables and parameters.

| symbol | meaning |
| --- | --- |
| Variables |  |
| $A$ | size of observation area |
| $N(A)$ | number of individuals in observation area $A$ |
| $S(A)$ | number of species in observation area $A$ |
| $P(A)$ | total productivity of individuals in observation area $A$ |
| $I(A)$ | invariability of productivity $P(A)$, i.e., the reciprocal of the squared coefficient of variation of $P(A)$ |
| Parameters |  |
| $\lambda_I$ | density of individuals within species range, i.e., for a given species the density of individuals belonging to this species in the range of this species; units of $\text{area}^{-1}$ |
| $\lambda_S$ | density of species range centers; units of $\text{area}^{-1}$ |
| $Q$ | size of species range |
| $\lambda_I Q$ | average number of individuals per species |
| $\lambda_S Q$ | for a given point of the landscape the average number of species ranges containing this point |
| $\lambda_I \lambda_S Q$ | total density of individuals, i.e., the density of individuals all species confounded; units of $\text{area}^{-1}$ |
| $m_p$ | mean productivity of an individual |
| $\sigma_p^2$ | variance of productivity of an individual |
| $\rho_{\text{intra}}(d)$ | correlation coefficient of the productivity fluctuations of two individuals belonging to the same species, i.e., intraspecific correlation |
| $\rho_{\text{inter}}(d)$ | correlation coefficient of the productivity fluctuations of two individuals belonging to different species, i.e., interspecific correlation |
| $\rho_0$ | correlation coefficient of the productivity fluctuations of two nearby individuals; specific to IAR-DD |
| $D_0$ | correlation area, i.e., for a given individual the area in which other individuals have productivity fluctuations correlated with this individual; specific to IAR-DD |

#### Spatial distribution

We consider a very large, spatially homogeneous landscape. First, we distribute species ranges over the landscape using a simple random process (a Poisson point process, see Appendix A in the Supporting Information). This random process is spatially homogeneous, that is, in any point of the landscape the number of species is the same on average. Species ranges do not change over time, and are assumed to be circular, with the same area for all species. Next, we distribute the individuals of the various species over their range, using a similar spatially homogeneous random process (see Appendix A). The positions of the plants do not change over time. The combination of the two random processes is characterized by three parameters: *Q*, the species range size; *λ*_*S*_, the spatial density of species ranges (more precisely, there are on average *λ*_*S*_*A* species range centers in an area *A* of the landscape); and *λ*_*I*_, the spatial density of individuals of a particular species within its range (in particular, each species has *λ*_*I*_*Q* individuals on average).

#### Temporal fluctuations

Plant productivity fluctuates through time. We assume that temporal mean and variance of these fluctuations are the same for all individuals. That is, denoting the productivity of individual *i* by *p*_*i*_,

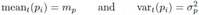

are the same for all individuals, and hence independent of species identity. To specify the spatial correlation structure of these fluctuations, we distinguish between intra- and interspecific correlations. The correlation coefficient of productivities *p*_*i*_ and *p*_*j*_ of two individuals *i* and *j* is

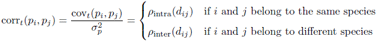

where *d*_*ij*_ is the distance between the two individuals.

### Species-area relationship (SAR)

The SAR relates the number of observed species *S*(*A*) and the observation area *A*. For simplicity, we only consider circular observation areas. The number of species *S*(*A*) can be easily computed for the homogeneous spatial structure of our model (see Appendix B). The derivation proceeds in two steps. First, we determine the set of species for which the range overlaps with the observation area. Second, for each of these species, we compute the probability that at least one individual is present in the region of overlap, which is a necessary condition to observe the species. The result of this computation can be expressed as a one-dimensional integral, see equation (B.2), which can be evaluated using numerical integration. The Supporting Information includes R code to compute the SAR.

As we will show in the results section, the SAR predicted by our model often consists of three phases. In Appendix B we derive a simple linear approximation for each phase, leading to a piecewise linear approximation for the entire SAR. We use the latter approximation, which is fully analytical, to systematically describe the parameter dependences of the SAR (see Table 2).

**Table 2.** Predictions of the piecewise linear approximation. The species-area relationship (SAR) and the two invariability-area relationships (IAR-DST and IAR-DD) are approximated by three line segments. The first one (column “first phase”) is linear in the area *A*, the second one (column “second phase”) is constant, and the third one (column “Cthird phase”) is again linear in the area *A*. The downward inflection point is approximated as the intersection of the first and second line segment; the corresponding area is given in column “downward”. The upward inflection point is approximated as the intersection of the second and third line segment; the corresponding area is given in column “upward”. Note that both IAR-DST and IAR-DD are proportional to the productivity invariability of a single plant, 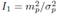.

|  | first phase | downward | second phase | upward | third phase |
| --- | --- | --- | --- | --- | --- |
| SAR | $\lambda_I \lambda_S Q \times A$ | $1/\lambda_I$ | $\lambda_S Q$ | $Q$ | $\lambda_S \times A$ |
| IAR-DST | $I_1 \lambda_I \lambda_S Q \times A$ | $1/\lambda_I$ | $I_1 \lambda_S Q$ | $Q$ | $I_1 \lambda_S \times A$ |
| IAR-DD | $I_1 \lambda_I \lambda_S Q \times A$ | $1/\rho_0 \lambda_I \lambda_S Q$ | $I_1/\rho_0$ | $2D_0$ | $I_1/2\rho_0 D_0 \times A$ |

### Invariability-area relationship (IAR)

Denoting the total primary productivity in observation area *A* by *P*(*A*), we are interested in the variability of *P*(*A*), which we quantify as the squared coefficient of variation,

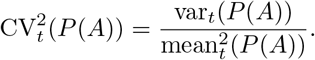

Low variability can be seen as an indicator of a stable ecosystem. Therefore, to obtain a proper stability measure, we take the reciprocal of variability, which is called invariability (Haegeman *et al.*, 2016; Wang *et al.*, 2017),

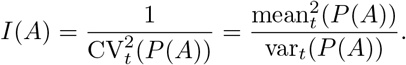

The IAR relates the invariability *I*(*A*) and the observation area *A*. In Appendix C we explain how this invariability can be computed for our model. For the mean in the numerator of *I*(*A*), we have

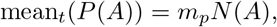

where *N*(*A*) is the number of individuals in *A*, which is proportional to the area *A*, see equation (C.2). For the variance in the denominator of *I*(*A*), we have

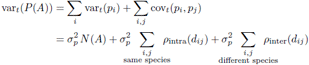

where the sums are over individuals *i* and *j* in the observation area *A*. In Appendix C we explain how the double sums in the last expression can be evaluated using Monte Carlo integration. The Supporting Information includes R code to compute the IAR.

#### Decorrelation by species turnover (IAR-DST)

While the previous analysis holds for arbitrary functions *ρ*_intra_(*d*) and *ρ*_inter_(*d*), in this paper we present results for two simple choices of these functions.

In the first case, we assume that individuals of the same species have perfectly correlated productivity fluctuations, and that individuals belonging to different species have independent fluctuations. This corresponds to setting

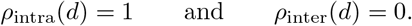

This admittedly extreme assumption could result from strong dispersal, so that entire species respond in unison to the environmental disturbances. In addition, plant species are assumed to have specific responses to the disturbances, so that productivity fluctuations are uncorrelated between species. Alternatively, the decorrelation between species can be generated by species interactions. For example, a combination of positive and negative species interactions, tending to increase and decrease species correlations, respectively, might cancel out species correlations on average. In any case, under this assumption the fluctuation correlations across space are governed by species differences. We call this decorrelation by species turnover (DST), and denote the corresponding invariability-area relationship by IAR-DST.

#### Decorrelation by distance (IAR-DD)

In the second case, we assume that the productivity correlations between any two individuals, whether they belong to the same species or not, only depend on the distance between the individuals (Wang *et al.*, 2017). This implies that intra- and interspecific correlations are identical, *ρ*_intra_(*d*) = *ρ*_inter_(*d*). Temporal correlations decay with distance, and one of the simplest functions describing this distance dependence is the exponential one (Bjørnstad & Falck, 2001; Liebhold *et al.*, 2004),

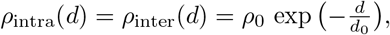

where *ρ*_0_ is the correlation between two nearby individuals, and *d*_0_ is the characteristic correlation length. For distances *d* well below *d*_0_ the correlation is equal to *ρ*_0_ and it vanishes for distances *d* well above *d*_0_. This assumption might be suitable for plant species whose productivity fluctuations reflect the variation of a dominant environmental variable, e.g., precipitation. Distance *d*_0_ would then correspond to the correlation length of the environmental disturbances. We call this assumption decorrelation by distance (DD), and denote the corresponding invariability-area relationship by IAR-DD.

As we will show, both the IAR-DD and the IAR-DST often have a triphasic shape. As for the SAR, these relationships can be approximated by a simple piecewise linear function (see Appendix C). We use this analytical approximation to study how the IAR depends on the model parameters (see Table 2).

## Results

### Species-area relationship (SAR)

Our model predicts a triphasic relationship between the number of species *S*(*A*) and the observation area *A* (on a log-log plot, Fig. 1). The slope switches from one at very small spatial scales (point A in Fig. 1), to a small value at intermediate scales (point B), and eventually to one again at large spatial scales (point C).

This shape can be characterized analytically using a piecewise linear approximation (Fig. 1 and Table 2). The transition between the first and second phase, which we call the downward inflection point, occurs at area 1*/λ*_*I*_. Recall that *λ*_*I*_ is the spatial density of individuals of a species within its range, so that 1*/λ*_*I*_ is the average area occupied by one individual in the species range. The transiton between the second and third phase, which we call the upward inflection point, occurs at area *Q*, which is the typical size of a species range. The number of species at intermediate scales (second phase), at which the piecewise linear approximation has zero slope, is equal to *λ*_*S*_*Q*, where *λ*_*S*_ is the spatial density of species range centers. The product *λ*_*S*_*Q* can be interpreted as the average number of species ranges present at an arbitrary point of the landscape. Note that while the zero-slope approximation is sufficient for our purpose, the piecewise linear approximation could be improved to get non-zero estimates for the exponent *z* of the power-law fit *S*(*A*) *α A*^*z*^.

The appearance of three phases can be intuitively understood as follows. First, note that any individual is surrounded by an area of size 1*/λ*_*I*_ on average in which no conspecific individual is present. Hence, if the observation area *A* is well below 1*/λ*_*I*_, there is only a small probability of observing two individuals of the same species. When increasing the observation area *A* (but still *A*⪡ 1*/λ*_*I*_), a newly observed individual belongs most probably to a species not previously observed (panel A of Fig. 1). As a consequence, the number of observed species *S*(*A*) increases linearly with the number of observed individuals, which is proportional to the observation area *A*. This explains why the SAR has slope one at small spatial scales.

Second, when the observation area *A* becomes comparable to 1*/λ*_*I*_, newly observed individuals often belong to already observed species, and the SAR exhibits a downward inflection. When further increasing the observation area (*A* ≫ 1*/λ*_*I*_), most of the species whose range overlaps with the observation area are effectively observed (that is, there is already a conspecific individual present in the region of overlap; see panel B). As long as the observation area *A* remains smaller than species range size *Q*, there are seldomly new species to be observed. This explains why the SAR plateaus at intermediate scales.

Third, when the observation area *A* becomes comparable to the species range size *Q*, new species ranges are appearing in the observation area, and the SAR exhibits an upward inflection. When further increasing the area (*A* ≫ *Q*), most observed species have their range centers included in the observation area (panel C). Hence, the number of observed species *S*(*A*) is approximately equal to the number of range centers in the observation area. This number, given by *λ*_*S*_*A*, is linear in *A*. This explains why the SAR has slope one at large spatial scales.

### Invariability-area relationship (IAR)

Our model also predicts a triphasic relationship between ecosystem invariability *I*(*A*) and observation area *A* (on a log-log plot, Fig. 2), both in the case of decorrelation by species turnover (IAR-DST) and in the case of decorrelation by distance (IAR-DD). The slope changes from one at very small spatial scales, to a small value at intermediate scales and again to one at large spatial scales.

**Figure 2.**
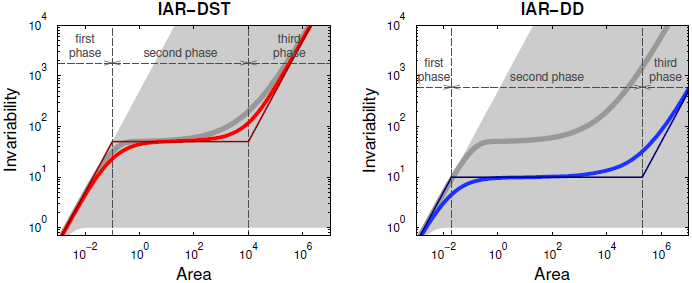
Our model predicts triphasic invariability-area relationships (IARs). We plot ecosystem invariability *I*(*A*) against observation area *A* under two decorrelation assumptions. Left panel: decorrelation by species turnover (IAR-DST). Exact model solution in thick red line and piecewise linear approximation in thin red line. Right panel: decorrelation by distance (IAR-DD). Exact model solutions in thick blue line and piecewise linear approximations in thin blue line. The SAR (thick grey line) is identical in the two panels. The shaded region indicates the set of possible IARs for a fixed configuration of individuals. For points on the upper boundary, all individuals have independent fluctuations; for points on the lower boundary, all individuals have perfectly correlated fluctuations. Parameter values: *λ*_*I*_ = 10, *λ*_*S*_ = 0.005, *Q* = 10^4^, *m*_*p*_ = 1, 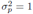, *ρ*_0_ = 0.1, *D*_0_ = 10^5^.

The three phases can be explicitly linked to the model parameters using a piecewise linear approximation (Fig. 2 and Table 2). This link allows us to gain insight into the underlying mechanisms of the IAR. In the IAR-DST, the downward and upward inflection points coincide with those of the SAR. Moreover, for each of the three phases, invariability is equal to the number of species multiplied by the invariability of a single plant. The latter invariability is equal to 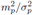, where *m*_*p*_ is the mean and 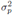 the variance of the productivity of a single individual. Hence, in the piecewise linear approximation, the entire IAR-DST coincides with the SAR up to the constant 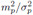. This result can be easily understood. As all individuals of the same species fluctuate in perfect synchrony, the invariability of the entire species, or of any part of that species, is equal to 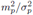 Invariability can only increase by including other species. Because different species fluctuate independently, invariability is additive in the number of species. Indeed, both the mean and the variance increase linearly with the number of species, so that invariability, which is equal to the squared mean divided by the variance, also increases linearly. This explains why the IAR-DST is proportional to the SAR.

In contrast, the IAR-DD is not directly linked to the SAR. The piecewise linear approximation indicates that the IAR-DD coincides with the IAR-DST in the first phase. However, at larger spatial scales, the IAR-DD strongly depends on two specific parameters: the correlation coefficient *ρ*_0_ between nearby individuals, and the area over which the correlations extend, 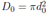 with *d*_0_ the correlation length. In particular, invariability of the second and third phases is inversely proportional to *ρ*_0_ and to *ρ*_0_*D*_0_, respectively. The piecewise linear approximation suggests a limited dependence on the spatial distribution of the ecosystem. Only the first phase depends on parameters *λ*_*S*_ and *Q*, and only so through the total density of individuals (equal to *λ*_*I*_*λ*_*S*_*Q*, see equation (B.1)). A more detailed analysis shows that there is a weak dependence on parameters *λ*_*S*_ and *Q*, due to the spatial clustering of individuals in species ranges (see Appendix C). This clustering strengthens the fluctuation correlations between individuals on average. As a result, invariability is slightly smaller than predicted by the piecewise linear approximation. Overall, in the case of decorrelation by distance, the spatial distribution of species, which entirely determines the SAR and the IAR-DST, has only a minor effect on the IAR-DD.

### Stability loss across spatial scales

As an application of our model, we explore how changes in the distribution of individuals and species affect the spatial scaling of ecosystem stability. We focus on two components of global change, species loss and habitat destruction, and describe their effects at multiple spatial scales. Obviously, we do not intend here to provide a realistic description of these complex phenomena and their consequences. Rather, we explore the variety of ways in which ecosystem stability can be affected from small to large scales.

We simulate different global change scenarios by varying species density *λ*_*S*_, individual density *λ*_*I*_ and species range size *Q*. We monitor the effects of these variations through the invariability-area relationship, assuming either decorrelation by species turnover (IAR-DST) or decorrelation by distance (IAR-DD). Recall that the SAR, which is independent of the decorrelation assumption, is directly proportional to the IAR-DST.

The first three scenarios deal with species loss. Scenario A simulates species loss by decreasing species density *λ*_*S*_ (Fig. 3A). As the remaining species are not affected, this causes the total density of individuals (equal to *λ*_*I*_*λ*_*S*_*Q*) to decrease as well. The IAR-DST decreases at all spatial scales, while the IAR-DD only decreases at the smallest scales and is unaffected at larger scales. This can be explained by recalling that the two IARs coincide in the first phase, and that the IAR-DD does not depend on the distribution of individuals and species in the second and third phases.

**Figure 3.**
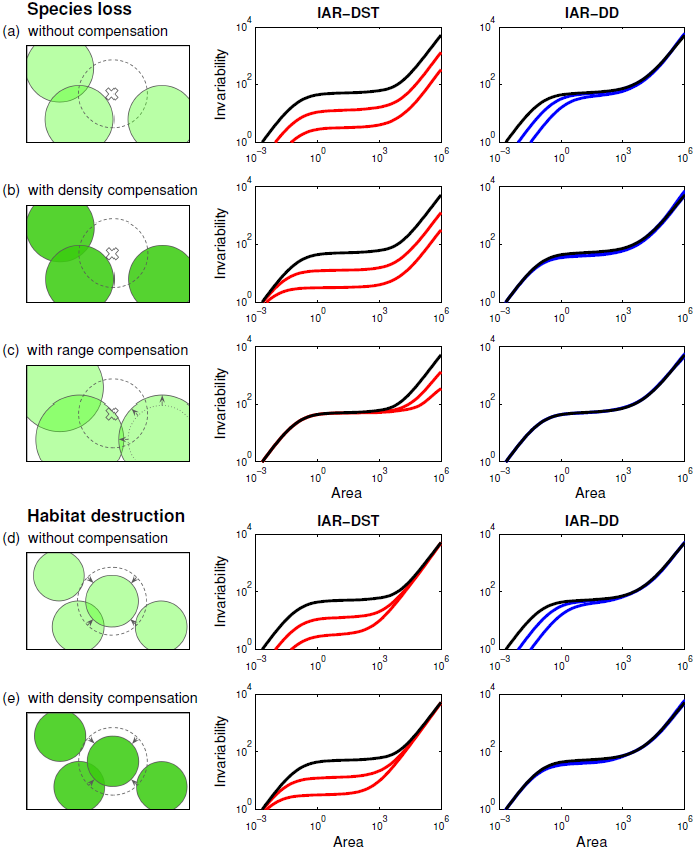
Simple scenarios of global change affect stability in various ways across spatial scales. We consider five scenarios: (a) species loss alone, (b) species loss associated with an increase of population density, (c) species loss associated with an increase of range size, (d) habitat destruction alone, and (e) habitat destruction associated with an increase of population density. Left column: illustration of simulated landscape. Middle column: IAR-DST. Right column: IAR-DD. Reference IARs for the initial landscape (black line) are the same across scenarios. Parameter values of the reference IARs are the same as in Fig. 2, except *ρ*_0_ = 1*/λ*_*S*_*Q* = 0.02 and *D*_0_ = *Q/*2 = 5 10^3^. Parameter changes are (a) *λ*_*S*_ *→ λ*_*S*_*/γ*, (b) *λ*_*S*_ *→ λ*_*S*_*/γ* and *λ*_*I*_ *→ γλ*_*I*_, (c) *λ*_*S*_ *→ λ*_*S*_*/γ* and *Q → γQ*, (d) *Q → Q/γ*, and (e) *Q → Q/γ* and *λ*_*I*_ *→ γλ*_*I*_, where the factor *γ* is equal to *γ* = 4 for the curve closest to the reference, and equal to *γ* = 16 for the curve furthest from the reference.

Scenario B simulates species loss by decreasing species density *λ*_*S*_ and simultaneously increasing individual density *λ*_*I*_ while keeping the total density of individuals (equal to *λ*_*I*_*λ*_*S*_*Q*) constant (Fig. 3B). This compensation could be due to competitive release, i.e., the extinction of a species creates the opportunity for its competitors to increase their density (Ives, 1995; Segre *et al.*, 2016). The IAR-DST decreases at all but the smallest scales, while the IAR-DD is not affected at all. The same explanation holds as in the previous scenario, with the addition that for both IARs the first phase is determined by the total density of individuals, which is constant in this scenario.

Scenario C simulates species loss by decreasing species density *λ*_*S*_ and simultaneously increasing species range size *Q* (Fig. 3C). This joint variation is such that the total density of individuals remains constant. This could occur if the extinction of species allows competing species to expand their range. In this scenario, the IAR-DST decreases only at large scales, where species density *λ*_*S*_ governs invariability. The IAR-DD does not change at any scale, as in the previous scenario.

The last two scenarios look at habitat destruction. Scenario D simulates habitat destruction by decreasing species range size *Q* (Fig. 3D). As all other parameters are kept constant, the total number of individuals decreases. For the two decorrelation assumptions, invariability decreases only at small scales. Indeed, for the IAR-DST, species range size *Q* does not affect the third phase, which is determined by species density *λ*_*S*_. For the IAR-DD, species range size *Q* does not affect the second and the third phases, which are determined by the correlation parameters *ρ*_0_ and *D*_0_.

Scenario E simulates habitat destruction by decreasing species range size *Q* and simultaneously increasing individual density *λ*_*I*_ (Fig. 3E), so that the total density of individuals remains constant. In this case habitat destruction reduces the space available per individual, but does not reduce the number of individuals. In comparison with the previous scenario, the IARs are not affected at the smallest scales, because the first phase is determined by the total density of individuals.

To sum up, despite its simplicity, our model predicts a rich variety of stability responses. Depending on the scenario considered, stability can be affected at a narrow or broad range of spatial scales, and predominantly at small, intermediate or large scales.

## Discussion

We constructed a minimal model that is able to predict simultaneously the species-area relationship (SAR) and the invariability-area relationship (IAR). For both relationships, we obtained a triphasic curve, in qualitative agreement with empirical data (Rosenzweig, 1995; Storch *et al.*, 2012; Wang *et al.*, 2017). It is remarkable that a simple model like ours is able to reproduce these patterns. Recall, however, that our model is not mechanistic, in the sense that it is not built on basic ecological processes. Instead, its starting point is simple assumptions about the spatial distribution of individuals and species and their temporal fluctuations. We translated these assumptions into predictions for the SAR and the IAR, but we did not connect them to underlying mechanisms. As a consequence, our model does not allow us to directly infer the ecological drivers of the SAR and the IAR (but see below).

In our model randomness serves a pragmatic purpose, i.e., it allows us to circumvent the complexity inherent in spatially extended ecosystems. But this does not mean that our model is incompatible with models that explicitly describe this complexity. In particular, irrespectively of the model complexity, the SAR and the IAR are entirely determined by the spatial configuration of the ecosystem and its fluctuation patterns. That is, once this spatio-temporal structure is given, the specific model details no longer matter for the SAR and IAR predictions. The basic idea of our model is to directly generate random instances of this structure, hence bypassing the underlying ecological complexity. Also, note that this model randomness does not refute the importance of deterministic processes (see also Coleman *et al.*, 1982).

We used the model to investigate the links between the SAR and the IAR. To demarcate the range of possible outcomes, we considered two assumptions about the temporal correlations of the ecosystem fluctuations. In the first case, called decorrelation by species turnover (DST), we assumed that species differences determine spatial asynchrony. We described a tight correspondence between the IAR-DST and the SAR, such that any process affecting the SAR has very similar effects on the IAR-DST. In the second case, called decorrelation by distance (DD), we assumed that spatial separation governs the decorrelation between ecosystem parts. We found that the IAR-DD is largely independent of the SAR, such that processes affecting the SAR often leave the IAR-DD unchanged. Nevertheless, despite this fundamental difference between the IAR-DST and the IAR-DD, there is a range of parameter values for which the shapes of the IAR-DD and the IAR-DST are similar. Indeed, the piecewise linear approximation indicates that a close match is obtained for *ρ*_0_ ≈ 1/*λ*_*S*_*Q* and *D*_0_ ≈ *Q*/2 (see Table 2). Hence, the same IAR can hide very different underlying processes.

We explored how simple scenarios of species loss and habitat destruction affect the IAR under the two decorrelation assumptions. We described a variety of stability responses, differing at small, intermediate and large spatial scales (Fig. 3). This emphasize the importance of investigating stability simultaneously at multiple scales. In particular, stability results at the local scale should not be extrapolated blindly, as this would represent a risk of underestimating (e.g., scenarios B and C under assumption DST) or overestimating (e.g., scenario D under assumptions DD and DST) large-scale stability. Note that the stability losses range over an order of magnitude (recall that invariability is represented on a logarithmic scale). We also found sharp differences between the response of the IAR-DD and that of the IAR-DST (e.g., in scenario B), even though the two IAR variants coincide for the reference scenario (black line in Fig. 3). This shows that the IAR by itself does not suffice to predict the stability effects of species loss.

Our model points at the missing information: the correlation functions *ρ*_intra_(*d*) and *ρ*_inter_(*d*). The more they differ, the larger the role of biodiversity in ecosystem stability, and the stronger the impact of species loss on the IAR. We focused on two extreme cases, in which the difference between *ρ*_intra_(*d*) and *ρ*_inter_(*d*) is either maximal (assumption DST) or zero (assumption DD). In real ecosystems, the correlation functions are somewhere intermediate between these extremes. It would be interesting to empirically evaluate *ρ*_intra_(*d*) and *ρ*_inter_(*d*). This should be possible from species-level community data consisting of time series at multiple spatial locations. This analysis would enable us to determine to what extent species diversity contributes to asynchrony in the ecosystem, and hence to its invariability. The answer, which most probably will depend on spatial scale, would be an indicator of the ecosystem’s vulnerability to species loss, regardless of the detailed processes governing the ecosystem.

Apart from the specific choices for the correlation functions, our model is built on several other simplifying assumptions. In particular, we assumed a strong degree of spatial homogeneity and species symmetry. Because the theoretical SAR and IAR are defined as averages over the landscape (e.g., the number of species *S*(*A*) is the average over all circular areas of size *A*), we do not expect that relaxing these assumptions will fundamentally modify our results. For example, if range sizes are allowed to differ between species, it would suffice to reinterpret parameter *Q* as the average range size (Allen & White, 2003). This need not hold for range size distributions with a long tail (e.g., a power-law distribution), which might also affect the slope of the SAR and the IAR at large spatial scales (see also the power-law decay of correlations considered by Wang *et al.*, 2017). Other model parameters can be made spatially heterogeneous and species dependent without qualitatively affecting the results.

For concreteness, we formulated the model in terms of plant productivity. However, our approach can be applied to other ecosystem properties and other taxonomic groups. For example, when considering mobile rather than sessile organisms, the movement of individuals introduces an additional contribution to ecosystem variability. A preliminary analysis shows that the corresponding IARs are similar to those studied in the paper. Also, the model interpretation given in the paper can be adapted to other levels of organization. For example, we assumed that individuals are the basic fluctuating entities, but in other contexts it might be more relevant to consider populations and their abundance fluctuations. Similarly, we assumed that species are the main entities that control decorrelation (at least under the DST assumption), but this role could also be played by functional groups, for example. In each of these interpretations, the effect of diversity on stability across spatial scales depends on the relationship between appropriately redefined correlation functions *ρ*_intra_(*d*) and *ρ*_inter_(*d*).

In summary, although not mechanistic, our modelling approach can accommodate a wide range of spatially structured ecosystems. Clearly, the next step is to connect this framework with ecological mechanisms. This would allow us to explain the driving parameters of our model, in particular *ρ*_intra_(*d*) and *ρ*_inter_(*d*), in terms of ecological processes, such as habitat selection, resistance to environmental disturbances and dispersal (for a similar approach in a metacommunty setting, see Wang & Loreau, 2016). Moreover, we would be able to mechanistically construct global change scenarios, rather than postulating them, as we were forced to do in this paper. Indeed, species loss and habitat destruction might affect various parameters of our model (e.g., *m*_*p*_, 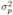 and the correlation functions), and more detailed models are needed to identify these multiple and interdependent effects (see Rybicki & Hanski, 2013 and Matias *et al.*, 2014 for analogous work on the SAR). Importantly, even for these more complex models, invariability is eventually determined by the spatial correlations of the fluctuating entities. Therefore, our approach can provide an integrative perspective on the spatial scaling of ecosystem stability and its link with biodiversity.

